# Within-host density and duration of colonization of multidrug-resistant Enterobacterales acquired during travel to the tropics

**DOI:** 10.1101/2023.03.03.530937

**Authors:** Olivier Cotto, Laurence Armand-Lefèvre, Sophie Matheron, Etienne Ruppé, François Blanquart, the VOYAG-R study group

**Author notes:** The VOYAG-R study group: Antoine Andremont, Laurence Armand-Lefèvre, Olivier Bouchaud, Yacine Boussadia, Pauline Campa, Bruno Coignard, Paul-Henri Consigny, Assiya El Mniai, Marina Esposito-Farèse, Candice Estellat, Pierre-Marie Girard, Catherine Goujon, Isabelle Hoffmann, Guillaume Le Loup, Jean-Christophe Lucet, Sophie Matheron, Nabila Moussa, Marion Perrier, Gilles Pialoux, Pascal Ralaimazava, Etienne Ruppé, Daniel Vittecoq, Ingrid Wieder and Benjamin Wyplosz.

## Abstract

Although it seems obvious that antibiotic use promotes antibiotic resistance, the processes underlying the changes in prevalence of MRE in European communities are still poorly understood. Information on how the within-host bacterial density varies after acquisition of a resistant strain and in the absence of antibiotic selection is key to our ability to further understand, model and manage resistance in the community. Empirical studies on the within-host dynamics following colonization by MRE are very scarce. Here, we study the within-host strain dynamics in healthy travelers colonized with MRE upon their return from tropical regions. Densities of both sensitive and resistant strains are stable for a few months, until resistant strains are abruptly cleared. Multivariate survival analysis further revealed that MRE acquired from Asia and carried at larger densities persist for longer. These dynamics do not support the classically assumed slow competitive exclusion of MRE. Rather, MRE and sensitive strains coexist in apparent equilibrium for months, with MRE representing about 0.1% of total Enterobacterales, before MRE are abruptly cleared. These results inform potential therapeutic strategies to clear MRE and epidemiological models of antibiotic resistance.

## Introduction

Multidrug-resistant Enterobacterales (MRE) pose major public health issues. Common types of MRE include Enterobacterales (EB) producing extended-spectrum β-lactamase (ESBL), plasmid encoded AmpC-type cephalosporinase and/or carbapenemase. ESBL-producing *Escherichia coli* strains represent the largest fraction of MRE in the community [1,2]. MRE can degrade most β-lactam antibiotics, which limits treatment alternatives in the cases of infection. The prevalence of MRE in developed countries has increased in the last two decades [2,3], but seems to have stabilized at 5-15% prevalence since around 2010 in some European countries [4,5]. Although it seems obvious that antibiotic use promotes antibiotic resistance, the processes underlying the changes in prevalence of MRE in European communities are still poorly understood [6,7].

Information on how the within-host bacterial density varies after acquisition of a resistant strain and in the absence of antibiotic selection is key to our ability to further understand, model and manage resistance in the community. Indeed, within-host dynamics determine the shedding of the resistant strain and potential onward transmission [8]. Understanding within-host dynamics can also guide the design of strategies to eliminate resistant strains. For example, the elimination of MRE strains by competition with ingested sensitive strains is envisioned to limit resistance in farm animals [9] and humans [10]. At the epidemiological level, recent mathematical modelling suggests that the emerging epidemiological dynamics of resistant and sensitive strains in the community and the equilibrium frequency of resistance critically depend on within-host dynamics [11]. Precisely, long-term coexistence of resistant and sensitive strains within the host after treatment could help maintain a moderate stable prevalence of MRE at the population scale. This result relies on the slow dynamics of competitive exclusion of the resistant strain by a sensitive strain in untreated hosts, occurring over several months.

Very few studies provide insights on the patterns of within host dynamics of resistant and sensitive strains *in vivo*. Indeed, longitudinal surveys of strain density during and after antibiotic treatment are challenging to implement. An experimental study of 20 piglets under 5-days ciprofloxacin treatment showed that the density of resistant strains increased over the course of treatment and shortly after (days 0-7), then briefly declined (days 7-12), presumably because of the competition with sensitive strains; from day 12 onward to day 27, no decline in the density of resistant strains was noticeable: they remained at a detectable density of approximately 1% of total Enterobacteriaceae [12]. A study of 133 patients longitudinally followed during hospital stays (lasting 15 days on average) evidenced a clear signal of selection for bacteria carrying CTX-M enzymes during 3^rd^ generation cephalosporin treatment. The study suggested that resistant strains are then slowly outcompeted by sensitive strains over relatively short time scales of 30 days [13]. However, in that study it is not clear what is the strength of the evidence for the declining density of resistant strains after treatment; the absolute value of the extinction threshold is tuned to match the median time to extinction of 30 days inferred in a separate study of returning travellers [8]; and the individual trajectories of resistance density in the 19 untreated individuals did not present a decline (Figure 1 right panel in [13]). We will come back to this study in more details and in comparison with our results in the discussion. To date, no other study investigated the change in resistant strain density until clearance in untreated hosts. More data analyses are required for further understanding of the within-host dynamics of resistant strains, especially over longer time scales and in healthy subjects.

**Figure 1:**
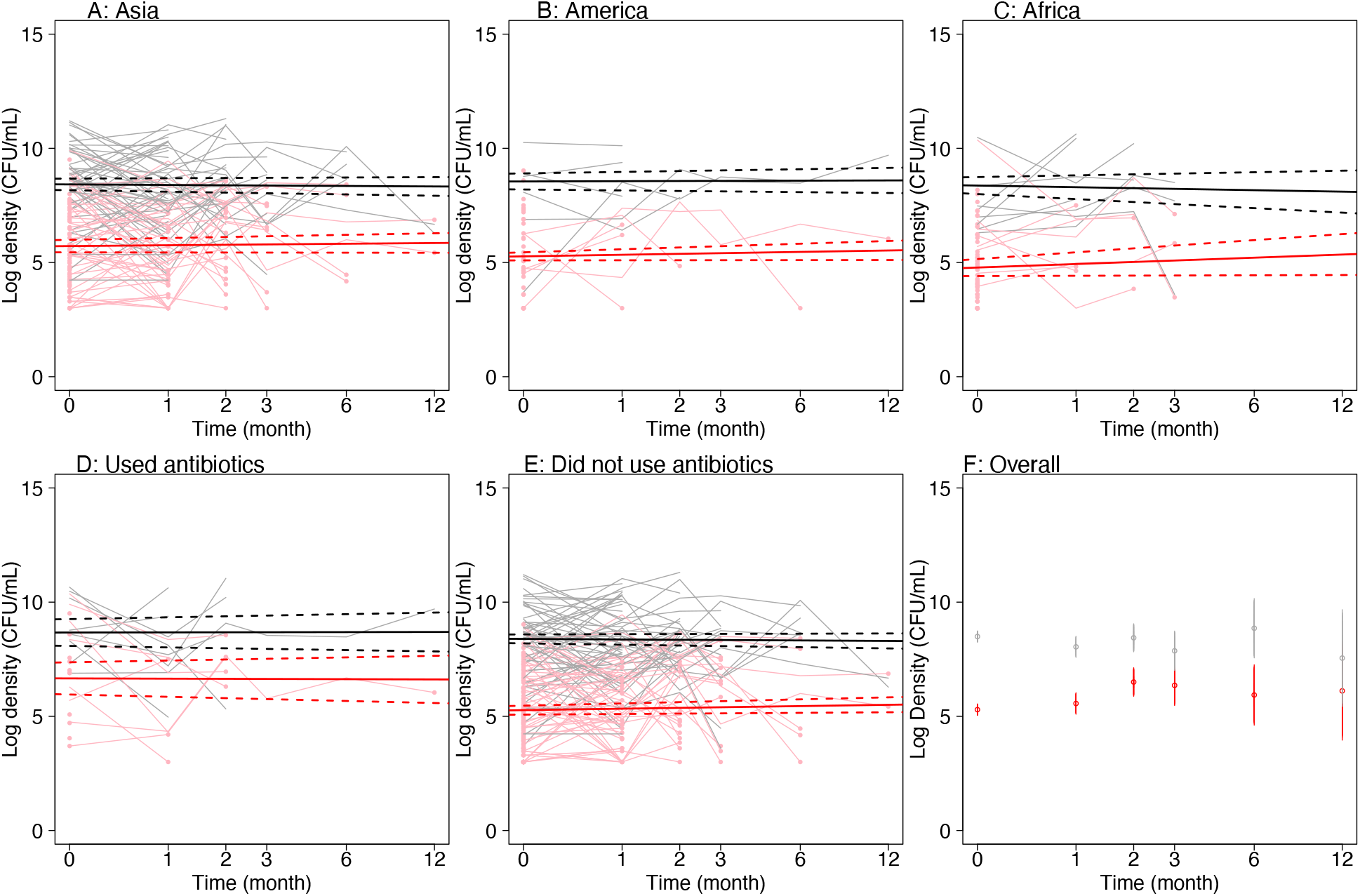
Density of bacteria as a function of time. Red and black correspond to MRE and EB densities respectively. Individual time-trajectories in bacterial densities are represented in light hue. Dots at the end of an individual MRE trajectory materialize the last time MRE were detected (single dots correspond to individuals who cleared their MRE during the first month). Thick solid lines represent linear model predictions and dashed lines represent the 95% C.I. A, B and C: individuals returned from each of the three travel regions. D and E, individuals who did and did not use antibiotics during travel, respectively. F: Mean densities as a function of time. Log is on base 10.

In the present study, we take advantage of data from a previously published monthly survey of both total EB and MRE density in a cohort of healthy travelers returned from tropical regions with MRE carriage [14]. Previous analyses of these data showed that, at the population level, short MRE carriage was associated with low MRE density relative to the total EB density. At the individual scale, the within-host dynamics of MRE density following travel has, however, not been investigated. In particular, the rate of decline of the density of resistant strains and the distribution of the time to clearance of these strains after return from travel has not been described.

Here, to get insights on the within-host MRE dynamics, we extend previous analyses from Ruppé et al. [14] by focusing on MRE dynamics at the individual level. We investigate both the dynamics of MRE density and the dynamics of total EB, from which we can deduce the density of sensitive EB strains. We discuss the implications of these results for our understanding of the mechanisms driving the within-host dynamics of resistant and sensitive strains after treatment, for the development of between-host models of resistance evolution, and for new approaches to eliminate MRE from the gut with probiotics.

## Methods

### Description of the study

Healthy individuals planning to travel to the three main tropical regions (sub-Saharan Africa, Latin America including the Caribbean, and Asia) volunteered to complete a basic questionnaire, provide a stool sample prior to travel and regular stool samples after their return. These samples were used to verify that no MRE was carried before travel and monitor the presence of MRE and the density of MRE over time after travel [14]. The questionnaire included information on malaria prophylaxis with doxycycline and the type of antibiotic used during travel, if any.

Only individuals who did not carry MRE before travel were included in the study. Participating individuals then provided a stool sample within a week after their return from travel. All samples were screened for MRE. The 292 individuals who were carrying MRE at return were asked to provide samples 1, 2, 3, 6 and 12 months later or until no MRE were detected. Individuals who did not carry MRE at return were not considered for follow up. For each stool sample, the total density of EB and the density of MRE (in CFU/mL) were quantified at reception of the samples. In parallel, an enrichment step allowed to detect whether MRE occurred at very low density [14]. Colony-forming units were all tested and were considered to be the same clone when they were of the same species, had the same morphotype and had the same antibiotic resistance profile. EB species were not identified, but most of the MRE were *E. coli* (93.3 %, see [15]). The details of the analysis of the stool samples can be found in Ruppé et al. [14]. Importantly, MRE were searched on fresh stool samples (not rectal swabs, self-collected at the traveler’s home and promptly shipped to the bacteriology laboratory of Bichat-Claude Bernard Hospital in Paris, France) using enrichment steps which increased the sensitivity for MRE detection [15]. This enabled detection of MRE at a density above 1000 CFU/mL. The details of the microbiological processing of the samples can be found in Ruppé et al. [14].

Previous analyses identified the factors associated with MRE acquisition in this cohort of travelers. In particular, the region of travel and the use of β-lactam antibiotics during travel had the strongest effects on the probability to carry MRE after return [14]. Here, we extend the investigation to the within-host dynamics of MRE density in a six-month period following return from travel. To this aim, we analyzed the density of both EB and MRE as a function of time for each individual in the survey and conducted a survival analysis of MRE strains following travel.

### Statistics

All analyses were performed using the software R [16]. The density data were log_10_-transformed. We visually verified that the transformed densities followed approximately a normal distribution. The relative MRE density was defined as the difference between the total log-density of EB and the log-density of MRE. The analyses were performed on the subset of 224 (among 292) individuals who acquired MRE during travel, who pursued the study until no MRE were detected in their stools (i.e. not lost to follow up), and for whom the MRE carried were *E. coli*. In total, the analyses were conducted on 353 samples (224, 64, 36, 18, 8, 3 at each sampling time – see above, respectively). A flowchart of the survey is provided in Figure 1 of [14]. The details of the statistical analysis are available in Supplementary Material as a R-markdown document.

Among the 224 individuals who acquired MRE during travel and who were kept in the analysis, 108 traveled to Asia, 70 to Africa, and 46 to America. A total of 60 individuals used antibiotics during travel (26 in Asia, 22 in Africa and 12 in America). Four types of antibiotics were used: doxycycline (here used for malaria prophylaxis only), nifuroxazide, β-lactam and fluoroquinolone. Preliminary statistical analyses showed that doxycycline and nifuroxazide did not have an effect on the density of MRE (see Supp. Mat.). These antibiotics were excluded from the remaining analysis. For simplicity, further use of the denomination antibiotics will refer to both β-lactam and fluoroquinolone.

First we investigated the change in density of EB and MRE over time using mixed linear models with individual identity as a random effect on the intercept (R-package lme4, [17]). We did not have enough data points to support models with an individual random effect on the slope. All the analyses were multivariate. Model selection was performed by first constructing the full model, with all potentially relevant explaining variables, and then building all possible nested models. The AIC criterion was used to select the best (i.e., most parsimonious) model where likelihood ratio tests were performed to obtain p-values associated with the remaining explaining variables.

Second, we used standard survival analyses (R-package survival, [18]) to investigate the rate of MRE clearance as a function of time since return from travel. We conducted both semi-parametric (Cox-models, Appendix 1) and parametric analyses. We fitted fully parametric models of clearance separately for each travel region with relative MRE density and the use of antibiotics during travel as explanatory variables. Therefore, the best model for each region reflects the persistence of the MRE strains acquired in this region and its dependence on MRE density and use of antibiotics. For parametric models, we compared an exponential hazard model, which assume that the instantaneous clearance hazard does not vary with time, with models of changing clearance rate with time such as the Weibull model and the log-logistic model. Parametric models provide full information on clearance rate as a function of time, contrary to semi-parametric (Cox) models.

## Results

### Density of MRE as a function of time

Our main result is that the within-host density of MRE did not vary with time after return from travel (Figure 1). Indeed, we did not find evidence that the density of MRE (conditional on MRE carriage) change with time after travel (*β*_*time*_ = 0.0509 [−0.067; 0.17] CFU/mL per month, 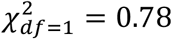, *N* = 353, *p* = .37, Figure 1). To investigate whether this result holds even for the subset of short-term MRE carriers, we further restricted the analysis to the hosts who cleared their MRE in the second month after their return (the mean half-time before clearance is between 1 and 2.5 months depending on the travel region, see below). These hosts provided two samples containing MRE, within a week after return and one month after return. In these short-term carriers, we also found that both absolute (*β*_*time*_ = −0.18 [−0.91; 0.54] CFU/mL per month, 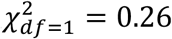, *N* = 56, *p* = .61) and relative MRE density (*β*_*time*_ = −0.22 [−1.03; 0.579] CFU/mL per month, 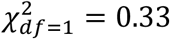, *N* = 56, *p* = .57) did not change significantly with time. In other words, in short-term carriers, MRE density did not vary during the first month, and MRE was cleared during the second month after return.

With a simulation study, we verified the power of our analysis to detect a within-host decline in MRE density and the accuracy of parameter inference. In this study we used the between-host variance and error parameters inferred from the linear model (Supplementary Figure S1). We found that the size of our data set and the level of error in density measures were sufficient to ensure a 100% power to detect a decline in density and a good accuracy of parameter estimates for relevant values of the within-host decline. The power dropped to 50% only for very slow declines in density of −0.1 per month: this slow decline would imply that MRE density falls to undetectable values on average after more than four years.

We further found in our data that the density of MRE increased with the number of distinct MRE strains (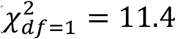, *N* = 353, *p* = .0007, *β*_nb.strains_ = +0.29 [0.10; 0.42] log_10_ CFU/mL per strain). The mean number of MRE strains upon return was 1.8 (median: 1 [1-8]). Moreover, consistent with previous results, we found that the travel region (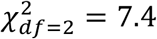, *N* = 353, *p* = .025) and the use of antibiotics (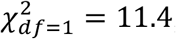, *N* = 353, *p* = .0007) contributed to the variation in MRE density. In particular, travelers returning from Asia had a larger density of MRE than travelers returning from other regions (*β*_Asia_ = 0.63 [0.19; 1.09] log_10_ CFU/mL and Figure 1) and travelers exposed to antibiotics had a larger density of MRE than those who were not (*β*_atb_ = 1.19 [0.41; 1.72] log_10_ CFU/mL, and Figure 1).

For the total EB density, we found that the null model (with the intercept only) performed best, such that none of the potential explaining factors significantly contributed to the observed variation (Figure 1). Consequently, the factors contributing to the variation in relative MRE density are similar to those explaining the variation in absolute MRE density (as presented above, see Supp. Mat.).

Among all EB carried by an individual, only a small fraction of cells was resistant. For each sample, we compared the likelihood of a model assuming that the total number of EB and MRE per sample followed different Poisson distributions with that of a model assuming that both numbers per sample followed the same Poisson distribution. In all cases, the best model suggested total EB density was significantly larger than MRE density (ΔAIC >> 2). On average, MRE represented 0.1% of the total EB population (Figure 1).

### Clearance rate of MRE strains after travel

Even though the density of MRE did not vary with time, the proportion of individuals carrying MRE decreased with time as MRE were cleared. We characterized the instantaneous clearance rate of MRE as a function of time. Preliminary Cox survival analyses (Appendix 1) indicated that both the use of antibiotics and the relative density of MRE had an effect on the clearance rate, but not the number of MRE strains carried. However, since the proportional hazard hypothesis was not met for the region of travel (Supp. Mat), we develop below our main analysis based on parametric models.

With parametric models, we found that log-logistic clearance models best described MRE clearance as a function of time (Figure 2). The instantaneous clearance rate was unimodal, first increasing with time to reach a maximum, and then decreasing (scale parameter < 1). The peak clearance rate occurred at about 2.5, 1.5 and 1.7 months for traveler from Asia, Africa and America respectively. Overall, the log-logistic models fitted well the pattern of MRE clearance in the first three months after travel (especially for Asia), but failed to capture MRE persistence later on (Figure 2).

**Figure 2:**
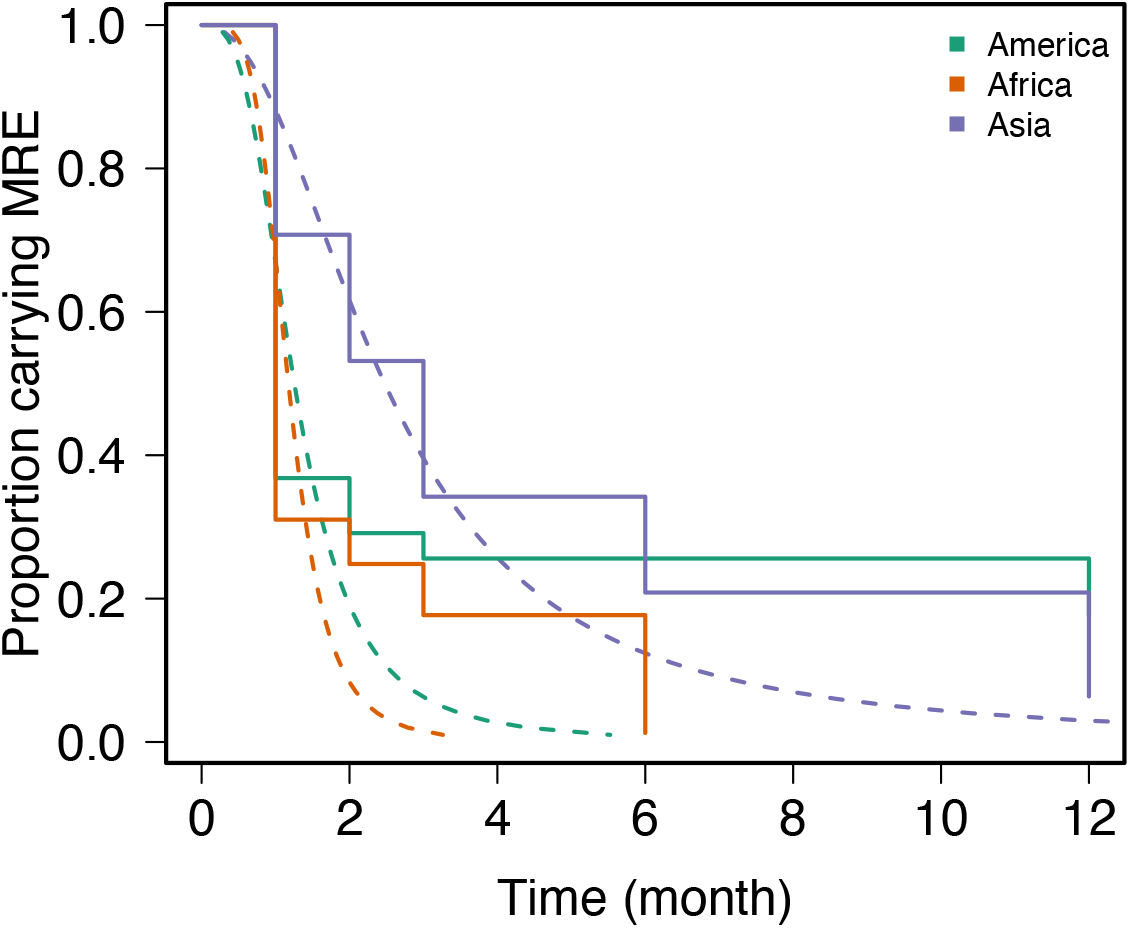
survival curves of MRE strains acquired in the different regions. Solid lines: Kaplan-Meier survival curves from the data; dashed lines: predictions from the log-logistic model. For the sake of illustration, the models include only the effect of the relative MRE density as a covariate. The effect of the use of antibiotics during travel are shown in Fig. S2.

MRE carried at large relative density persisted for longer. The effect of relative MRE density was strongest in travelers returning from Asia (a factor 1.18 [1.10-1.26] per log_10_ CFU) followed by Africa (1.04 [1.01-1.09] per log_10_ CFU) and Latin America (1.02[0.93-1.10] per log_10_ CFU).

The effects of the antibiotic treatment on the persistence of MRE strains depended on the travel region (see details in Supp. Mat.): the use of antibiotics during travel decreased the persistence time of MRE strains (after return) by a factor 0.59 [0.39-0.88] in travelers from Asia, but increased it by a factor 2.73 [1.51-4.95] and 1.59 [1.08-2.33] in travelers from Latin America and Africa, respectively.

According to the survival models, the half-life (time at which 50% of individuals have cleared MRE) was 2.47, 1.17 and 1.24 months in travelers returning from Asia, Africa, and America respectively, when carried at the median relative density. The equivalent figures for MRE carried at the lower 5% density were 1.5, 1.17 and 1.24 months in travelers returning from Asia, Africa and America respectively. For MRE carried at the higher 95% density, they were 3.64, 1.36 and 1.34 months.

## Discussion

Our study provides new insights on the within-host dynamics of travel-acquired MRE in the gut. Our first main result is that clearance occurs abruptly after a period of stable MRE density that can last several months (Figure 1 and Figure 3 scenario 1). This scenario contrasts with the classical expectation that clearance is the end-result of a slow process of competitive exclusion (Figure 3, scenario 2). Our second main result lies in the detailed characterization of MRE clearance as a function of time, travel region and within-host density. This reveals distinct dynamics for strains from different regions, MRE from Asia being the most persistent, and an effect of MRE density on persistence.

**Figure 3:**
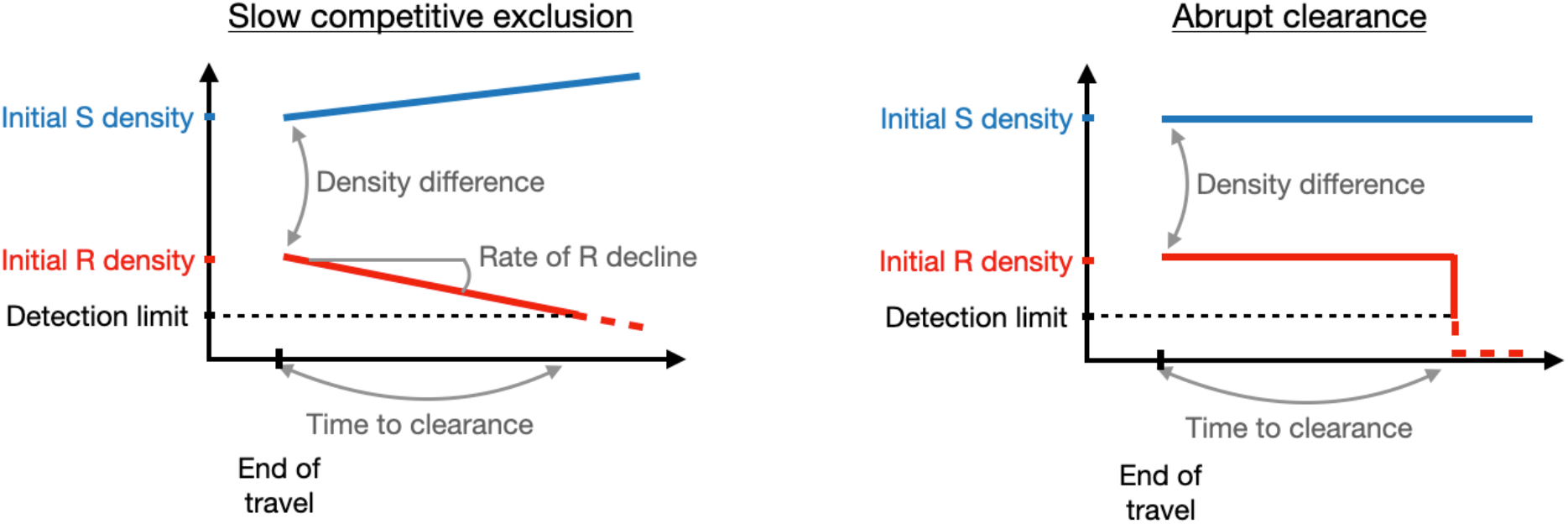
Alternative scenarios of MRE clearance. Left panel: after return from travel, MRE are slowly outcompeted by sensitive strains, until the density of MRE falls below the detection threshold. Right panel: The density of MRE at return from travel does not change with time, but MRE are eventually cleared. Our results favor this second scenario.

The mechanisms associated with MRE clearance are not known. We found that the clearance rate of MRE decreased with their density, suggesting that density-dependent mechanisms are involved in MRE clearance. Competitive exclusion of resistant strains by sensitive ones has been proposed as a candidate mechanism [11]. Indeed, resistance mechanisms often induce a fitness cost, which decrease competitivity in the absence of antibiotics [19]. Our results are not in contradiction with this view, but we showed that if competitive exclusion does occur, its effect on MRE strain density and the eventual clearance can occur as a brief event after a relatively long period of stable coexistence between resistant and sensitive strains.

How do our results compare to previous evidence? As explained in the introduction, the best previous study on longitudinally followed-up hospitalized individuals clearly demonstrates antibiotic selection on CTX-M-carrying bacteria, but does not give much details on the strength of the evidence for a decline in resistant strain density after treatment, and on the absolute value of this decline [13]. In an effort to understand this decline in greater details, we re-analyzed the dynamics of resistance in these data, focusing on never-treated individuals (N=19) and on treated individuals in untreated periods (N=51). We further stratified untreated periods into periods following 3^rd^ generation cephalosporin treatment (ceftriaxone, cefuroxime: C3G) or other treatments (Supplementary Figure S2). We found no decline of resistance in never-treated individuals (linear model, *β*_*time*_ = −0.0015 per day [−0.037, 0.034], p = 0.93), nor in individuals treated with non-C3G treatment (linear model, *β*_*time*_ = 0.00033 per day [−0.041, 0.042], p = 0.99). However, after C3G treatment, the density of resistance was elevated and there was evidence for a decline in density in the first few days post-treatment (linear model, *β*_*time*_ = −0.14 per day [−0.19, − 0.088], p = 2.6 10^−7^). These results from hospitalized individuals under C3G treatment are thus fully consistent with experimental results on piglets under ciprofloxacin treatment [12] and with our own results. Together this suggests biphasic dynamics of resistance post-treatment: first, over a few days, a rapid decline in resistance density as the sensitive strains re-establish and grow; second, over a few weeks to months, a stable equilibrium at which resistant strains persist at a small fraction of total density, before being eventually cleared.

We propose two mechanisms of note that could result in these clearance dynamics. Clearance may be caused either by the coexisting sensitive strains, or by displacement by new incoming sensitive strains. The first mechanism would require threshold effects on strain dynamics, as observed in other systems [20,21], to explain the sudden clearance after weeks or months of stability. The second mechanism requires that some combinations of strains stably coexist within hosts, but other incoming strains might very quickly disrupt the equilibrium. In any case, the coexistence over several months of resistant and sensitive strains, without change in density, poses the question of the mechanisms permitting the persistence of resistant strains despite their much lower density reflecting a fitness cost. This is a significant resistant subpopulation: 0.1% of the total density of Enterobacterales implies of the order of 100 millions resistant cells persisting in the gut for extended periods of time in apparent equilibrium with the sensitive strains, in spite of their cost. Resistance genes could be associated with genes enhancing persistence [22], increasing the mean time before colonization by a competing sensitive strain. Further research is needed to understand this intriguing long-term strain coexistence and the mechanisms of strain replacement.

Density-dependent mechanisms leading to MRE clearance could also occur at the scale of Enterobacterales species other than *E. coli*, or other members of the microbiota. Most (> 90%) MRE were identified as *E. coli*. Total Enterobacterales (the vast majority of which were sensitive) were not identified at the species level. In a similar protocol investigating stool samples of healthy individuals, *E. coli* account for around two thirds of total Enterobacterales. Moreover, a study from our group reported that a rapid MRE clearance (within one month after return) was associated with a high relative abundance of *Bifidobacterium* spp. and bacteria from the Clostridiales order in the gut [23]. Regardless of whether competition occurs at the intra- or interspecific scale, clearance is an abrupt process without prior decline in density.

The simulation study indicated that our longitudinal design had a 100% power to detect plausible fast and slow declines in MRE density, and good accuracy to estimate this decline. Of course, clearance must be preceded by a decline in MRE density, although the timescale of this decline was too fast to be observed. The analysis of MRE density focusing on the subset of travelers who lost their MRE between the first and second month indicates that density did not change during the first month, consistent with the overall analysis and suggesting the timescale of the decline is much faster than one month. Lastly, we did not have information on the use of antibiotics or health care after return from travel. Travelers following the survey were healthy, and the average rate of antibiotic consumption in the adult population low [24]. We expect that the use of antibiotics during the survey was sufficiently rare not to affect our overall conclusions.

Our survival analysis showed that the clearance rate is best described by log-logistic functions of time, with parameters depending on the travel area, on the density of MRE, and on whether travelers used antibiotics. Log-logistic clearance rates implies that the clearance rate increased with time to reach a peak before decreasing. The peak probability of clearance occurred between one and three months after return from travel, consistent with previous estimates of median MRE carriage time [14]. The variation of the clearance rate with time could reflect the diversity of MRE strains acquired during travel. Genomic analyses the 11 long-term carriers from the present data set showed that MRE strains vary in their persistence ability, with a single strain responsible of all carriage time exceeding three months [22]. At the population level, a diversity of MRE strains with variable persistence ability can result in an unimodal clearance rate as a function of time. The longer persistence of MRE acquired in Asia could be explained if persistent MRE strains are more likely to circulate in Asia than in other regions.

The timescale of MRE clearance has several implications. First, it informs the structure of mathematical models which aim to explain MRE prevalence at the community level. For example, Davies et al. [11] modeled a succession of states corresponding to a slow (several months) competitive replacement of resistant by sensitive strains. Our analysis shows that modeling competitive replacement as a slow process may not be appropriate. Second, ingested probiotics have been recently proposed to competitively replace MRE strains in the gut [10]. This strategy has further received some support from preliminary assessments in chickens [9,25]. Our results suggest that competitive exclusion by probiotic might occur in one or two ways. Along the first mechanism, competition between strains results in a period of stability over several months, followed by rapid clearance. Along the second mechanism, some combinations of MRE strains and probiotic strains would not result in clearance of MRE strains, while others would be highly effective and rapidly clear MRE. It would thus be important to distinguish between the two mechanisms. In future work, one way to do so would be to follow the within-host dynamics of *both* MRE and sensitive strains to model jointly their dynamics and turnover rates. We expect that our results will be of interest for future analyses of the role of competition to exclude MRE from the gut and for the development of models to understand and predict variation in MRE prevalence at the community level.

## Ethical Statement

The VOYAG-R study was approved by the Ile de France IV ethics committee on 14 November 2011. The study was observational and did not directly benefit the participants. French law requires that each participant sign a “nonrefusal” form. When MRE (including CPE) carriage was detected, the individual concerned was sent an information leaflet on MRE carriage, basic hygiene, and the need to mention this carriage when receiving medical care. Individuals with CPE carriage received a similar leaflet and were also contacted by the infection control practitioner of Bichat-Claude Bernard Hospital to ensure the information was correctly understood.

## Supplementary Figures

**Figure S1:**
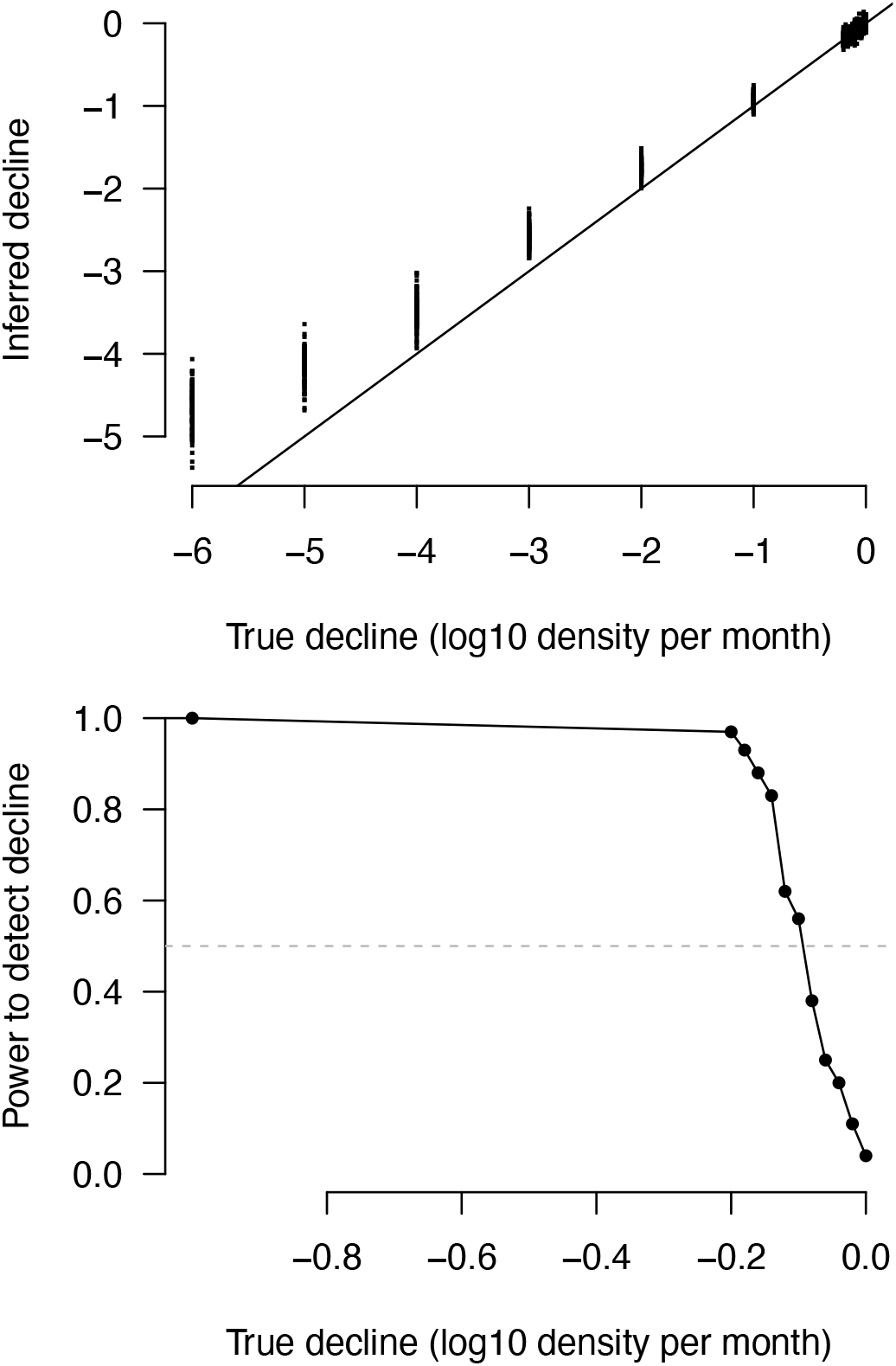
Analysis of the statistical power of a linear model with individual identity as a random effect (that we used) to detect a decline in MRE with time. The details of the analysis are provided in the main text (section “Density of MRE as a function of time”). 100 data sets were simulated for each rate of decline. The code is provided in supplementary material (R-markdown file). Upper panel: decline inferred by the linear model (dots) as a function of the true decline. Lower panel: frequency of detection of a significant decline (p-value < 0.05) as a function of the simulated MRE rate of decline.

**Figure S2:**
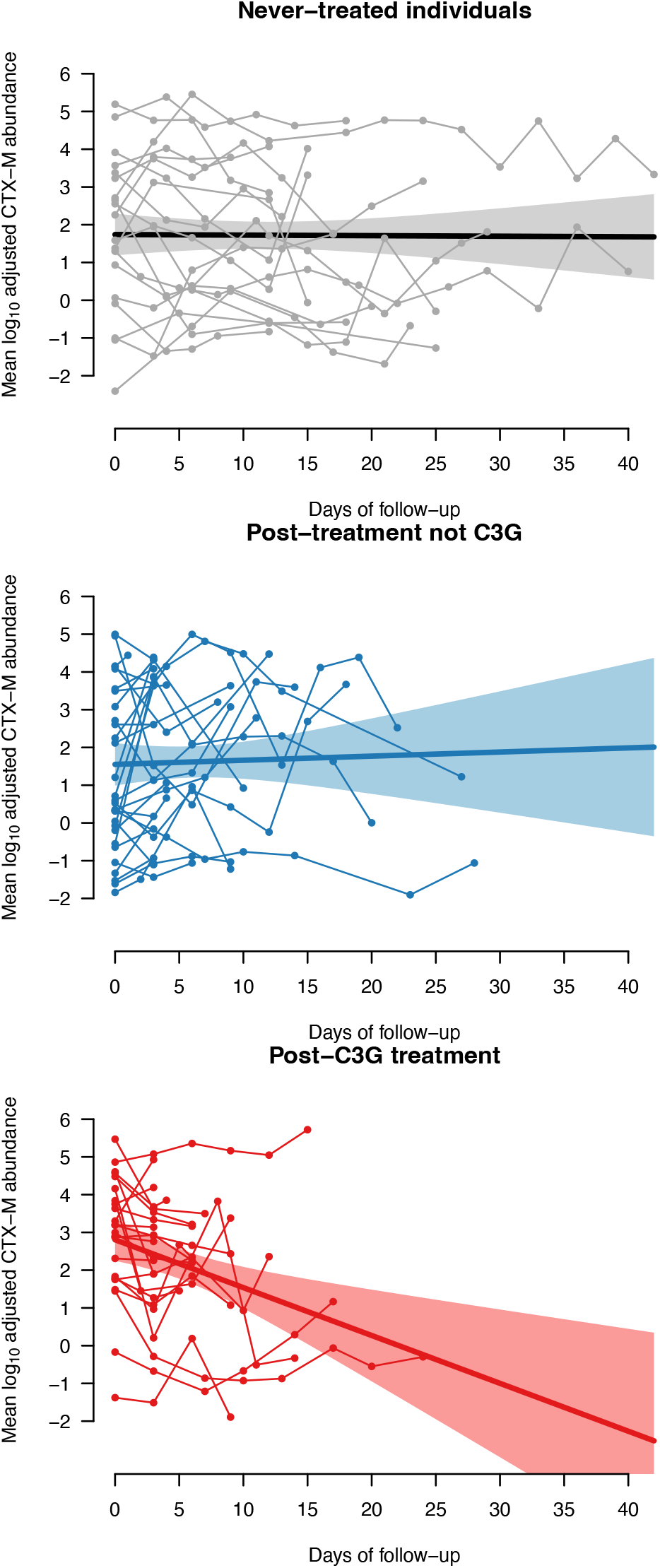
CTX-M abundance as a function of time in data from Niehus et al. [13]. A decline in CTX-M abundance is detected only following C3G treatment. Joined dots: per-individual CTX-M abundance as a function of day of follow up. Solid thick line: Linear model for the log_10_ CTX-M abundance as a function of the days of follow up (colored area: 95% CI).

